# An active model for the basilar membrane and the outer hair cells

**DOI:** 10.1101/2024.08.29.610286

**Authors:** Jorge Berger, Jacob Rubinstein

## Abstract

A model for the joint motion of the basilar membrane (BM) and the outer hair cells (OHC) in the cochlea is presented. The model consists of two one-dimensional mass distributions, one along the OHC and outer hair bundle (OHB) interface, and one along the BM. The motion of these masses is driven by the forces exerted on them by the elastic bodies connecting them and by the pressure difference in the fluids separated by the BM. The model includes a nonlinear motility of the OHC and its coupling with the vibrations of the BM. The model implies a Hopf bifurcation for the dynamical system governing the two coupled distributed oscillators. It is shown that when the system operates near the bifurcation point the BM motion is amplified up to a saturation level. The model provides very sharp frequency decomposition of the incident audio signal according to the place principle. It also acts as a powerful filter that distinguishes pure tones even in the presence of louder noisy background. In addition to simulations of the model, the unusual role played by the OHC friction is studied. Energy estimates are derived for the model functions.

## 1 Introduction

It is well-established [2], [13] that the cochlea functioning includes an active mechanism that amplifies the response of the basilar membrane (MB) and other components of the Organ of Corti (OoC) to an incident sound. The active element is responsible for a number of observed phenomena, including amplification, otoacoustic emission, sharpening the spectral decomposition of sound, saturation at large amplitude, noise filtering and more. In particular it is known that the energy source for the amplifier resides in the scala media, and it is activated via a network of ionic channels, triggered by the molecule prestin [5]. Several models were proposed for the mechanical nature of the active mechanism. From a mathematical point of view it is natural to assume that the fundamental dynamical behavior is a manifestation of a Hopf bifurcation [12], [13]. Indeed several authors used an ad-hoc canonical Hopf formulation to argue that this paradigm yields all the observed phenomena, e.g. [19], [4], [14].

In a previous publication [3] we considered a slice of the OoC. We wrote down Newton’s laws for many of its elements, including the Deiters cells (DC), the outer hair cells (OHC), the outer hair bundle (OHB) and more. Finally we wrote down approximate equations for the fluid motion in the endolymph that drives the inclination of the inner hair bundle (IHB). The motion of the BM entered our model as input for the OoC dynamics. One of the key features of our modular scheme is a model for the motility of the OHC. Specifically, we assumed that inclination of the OHB triggers contraction of the OHC. We showed that our model gives rise to a Hopf bifurcation, and through the chain of energy transfer from the BM to the IHB, it provides the desired effects of the cochlea active mechanism.

In the present paper we consider a complementary point of view. We concentrate on the BM and its coupling to the active OHC. The model is introduced in the next section. In section 3 we concentrate on the pure dynamical core of the model, ignoring the pressure difference across the BM. In section 4 we derive energy estimates for the full model. In section 5 we report on simulations of the model. We discuss our results and possible extensions of them in section 6. While the one-dimensional fluid-BM interaction model is well-known, and in fact was derived rigorously in [17] and [22], we briefly present a formal derivation of it in an appendix.

## 2 The model

A schematic view of our model is depicted in Figure 1. While the cochlea in general, and the OoC in particular, consists of an intricate network of mechanical components, we focus here on two of them where key signal processing activities take place: The OHC and the BM. The BM is modeled as a one dimensional elastic body. Its vibrations are driven by the pressure difference between the scala media on one side of it and the scala tympani on the other side. The reduction to one dimension was justified mathematically in [22]. Indeed the reduction is not uniformly valid, and particularly so near the position of maximal deflection, where the wavelength is comparable to the thickness of the compartments separated by the BM; however it was shown numerically to be in good agreement with the full hydromechanical model. The DCs are neglected here. The three rows of OHCs are modeled as a continuous one-dimensional elastic body. In practice we assume that there are two one-dimensional (1d) mass distributions: One along the BM and one along the OHC-OHB interface. The motion of these mass distributions is according to Newton’s laws, where the only unusual feature is the nonlinear motility of the OHC.

**Figure 1:**
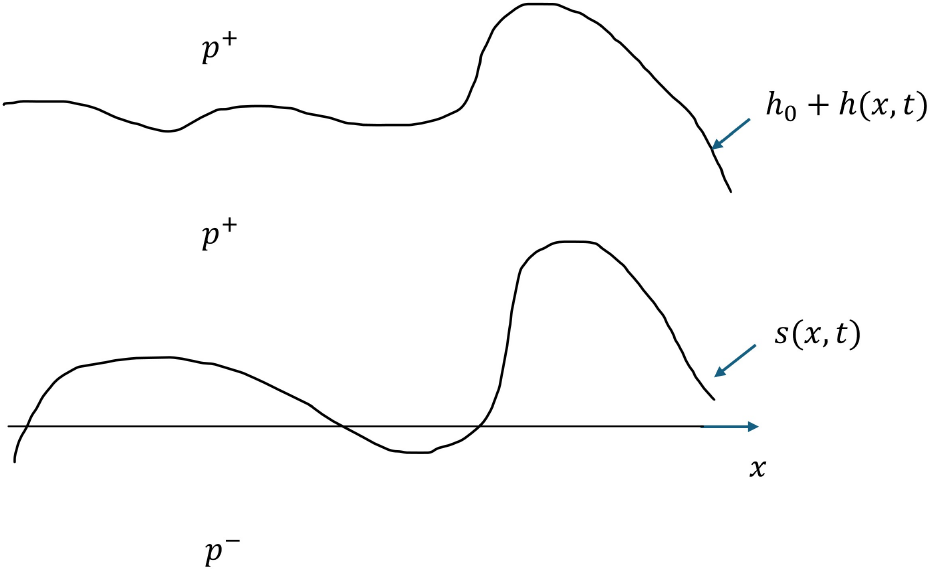
A sketch of the 1d model. The deflection of the BM with respect to its rest position (depicted as the *x*-axis) is denoted *s*(*x, t*), while *h*(*x, t*) is the vertical shift of the OHC-OHB interface relative to its rest position *h*_0_. The fluid pressures in the scala media and scala timpani are denoted *p*^+^ and *p*^−^, respectively.

### Units

We essentially use the same units as in [3]. Although the units there were selected, in part, in relation to the flow in the endolymph, that we do not consider here, it is convenient to adhere to them with a view towards a comprehensive OoC model in the future. The only exception is the horizontal length scale for the BM, taken as the stretched out length of the BM, that we select to be *L* = 30mm. Thus, time is measured in units of 10^−4^s, the vertical length unit is 10 µm, and the mass density unit is 10^−7^kg/m. The elastic constants *k*_*c*_, *k*_*d*_ below are in units of 10 *kg*/*ms*^2^, and the friction coefficients below are in units of 10^−3^kg/m s.

### The governing equations

For the pressure difference *p*(*x, t*) across the BM we use the one-dimensional model ([1], [17], Appendix A):

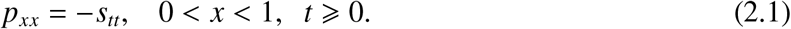

Here *s*(*x, t*) is the deflection of the BM, *x* denotes the location along the BM, and *t* denotes time. The system is driven by *p*(0, *t*) = *f* (*t*) the pressure difference at *x* = 0. We also assume *p*(1, *t*) = 0. The length of the uncoiled BM is normalized to 1. Neglecting the motion of the DC, the BM deflection is coupled to the motion of the OHC. The vertical shift of the OHC-OHB interface, relative to its rest position *h*_0_, is denoted *h*(*x, t*). Following [3] we assume that the coupled motion of the BM and OHC is activated by the forces applied by the continuum of springs connecting them and is described by

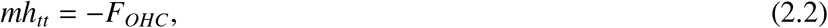

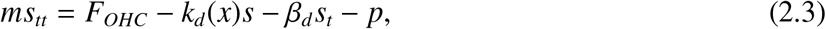

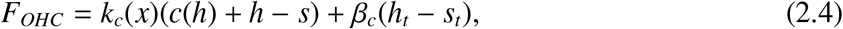

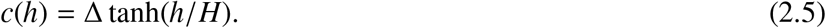

In particular we assume the motility of the OHC to depend on the deflection of the OHB via the nonlinear contraction term *c*(*h*) in equation (2.5). We take homogeneous initial condition for *h, s* and their first derivatives. We denote by *k*_*c*_ and *k*_*d*_ the (positive) elastic coefficients, the (positive) friction coefficients are *β*_*c*_, *β*_*d*_, The nonlinear term *c*(*h*) is saturated at *H* and Δ is a control parameter.

Notice that we assumed above that all the coefficients, except for *k*_*c*_(*x*) and *k*_*d*_(*x*), are *x*− independent. It is easy to generalize this but it would not affect the results qualitatively. We also take the mass densities of the BM and the OHC to be equal. This assumption is not essential and does not affect the results qualitatively.

## 3 The pure nonlinear oscillator

It is useful to study first the dynamical system (2.2)-(2.5) by itself at a specific location *x* and without the pressure difference term *p*. We thus look at the dynamical system

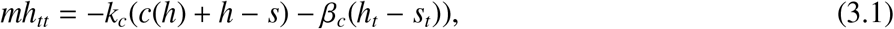

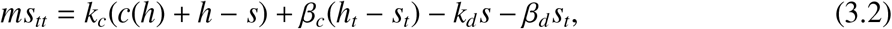

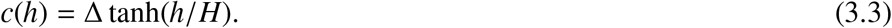

To study the stability of the oscillator (3.1)-(3.3) we convert it to a 4×4 first order system and linearize about *s* = *h* = 0:

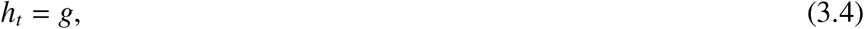

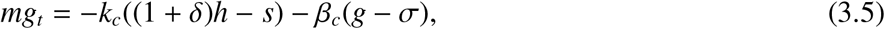

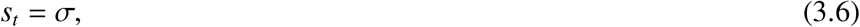

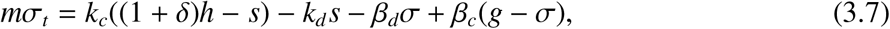

where we used *δ* := Δ/*H*.

Dividing the second and fourth equations by *m*, the stability of the system (3.4)-(3.7) is determined by the spectrum of the matrix

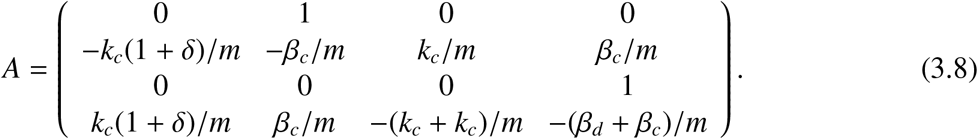

The matrix *A* has two pairs of complex conjugate eigenvalues. For small values of *δ* the real part of each pair is negative and the solution is at a stable mode only. However, when *δ* crosses a critical value denoted *δ*_*c*_, the real part of one eigenvalue pair becomes positive, while the imaginary component is nonzero, i.e. the system undergoes a Hopf bifurcation and an unstable mode emerges. For instance, when using the parameters *k*_*c*_ = 50, *k*_*d*_ = 400, *m* = 10, *β*_*c*_ = 43, *β*_*d*_ = 3, *H* = 0.0065 we obtain *δ*_*c*_ = 4.34. The critical frequency in this example, namely the complex component of that eigenvalue pair is *ω*_*c*_ = 4.5. Although not relevant to the cochlea functioning, we point out that increasing Δ much further, say to Δ = 0.8 drives the system back into decaying modes only.

**Remark:** To simplify the presentation we omit the parameter *m* in the sequel by rescaling the parameters *k*_*c*_, *k*_*d*_, *β*_*c*_, *β*_*d*_, Δ with respect to it.

### *Expansion near β*_*c*_ = *β*_*d*_ = 0

One of the interesting and illuminating features of this system is its behavior when the friction coefficients *β*_*c*_, *β*_*d*_ are very small. When both coefficients vanish the solution is purely oscillatory as we show below. What is unusual here is that when we keep *β*_*d*_ = 0 but slightly increase *β*_*c*_, then under a specific condition on the other coefficients appearing in the system (3.4)-(3.7), one pair of purely imaginary eigenvalues gains a *positive* real part; namely, the friction *β*_*c*_ destabilizes the dynamical system. These facts and more are expressed in the following

#### Proposition 1

The eigenvalues of the matrix *A* are purely imaginary when *β*_*c*_ = *β*_*d*_ = 0. Upon increasing *β*_*c*_ the dynamical system (3.4)-(3.7) undergoes a Hopf bifurcation if and only if 2*k*_*c*_*δ* > *k*_*d*_. When this condition holds, then in the linear regime the amplitude of the oscillations of *s* is larger than that of *h*. To justify this proposition we compute explicitly the eigenvalues λ(*β*_*c*_, *β*_*d*_) of *A*. A direct computation gives

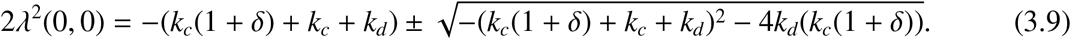

This formula implies at once that there are two pairs of purely imaginary conjugate eigenvalues. Next we perturb these eigenvalues by slightly increasing *β*_*c*_. Writing 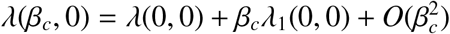 the leading order term is

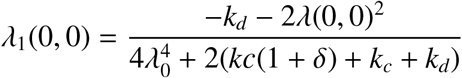

Using formula (3.9) we obtain after a little calculation that, for the eigenvalues associated with the plus sign in (3.9), λ_1_(0, 0) is a real number and furthermore

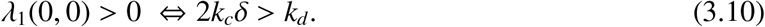

Finally, denoting by *u* = (*u*_1_, *u*_2_, *u*_3_, *u*_4_) the eigenvector associated with the bifurcating eigenvalue corresponding to (*h, g, s, σ*), we obtain after another little calculation that the condition 2*k*_*c*_*δ* > *k*_*d*_ implies |*u*_3_| > |*u*_1_|. We comment that when *β*_*d*_ > 0 there is also an upper bound on the range of *δ* for which the real part of λ is positive.

### Amplification and saturation of the pure oscillator

To see the response of the pure oscillator model to applied force at different amplitudes we introduced into the right hand side of equation (3.2) a driving force *χ*(*t*). The system (3.1)-(3.2) was then simulated under

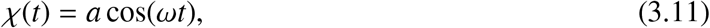

for four values of the amplitude *a* = 10^−2^, 10^−3^, 10^−4^, 10^−5^. We used the mechanical and saturation parameters *k*_*c*_ = 5, *k*_*d*_ = 40, *β*_*c*_ = 4.3, *β*_*d*_ = 0.3, *H* = 0.0065 and the bifurcation parameter Δ = 0.0282. To recall, the critical frequency in this example is *ω*_*c*_ = 4.5. In Figure 2 we depict the (logarithm of) maximal value of *s*(*t*)/*a* for the interval 4 < *ω* < 5 and for the four values of the driving amplitude *a*. The top graph is for *a* = 10^−5^ and the other graphs are monotone with increasing values of *a*. This result demonstrates that the pure oscillator provides both the amplification of the driving force and the saturation at high amplitudes. Notice also the slight shift to the left of the maximum of the graph for *a* = 10^−3^.

**Figure 2:**
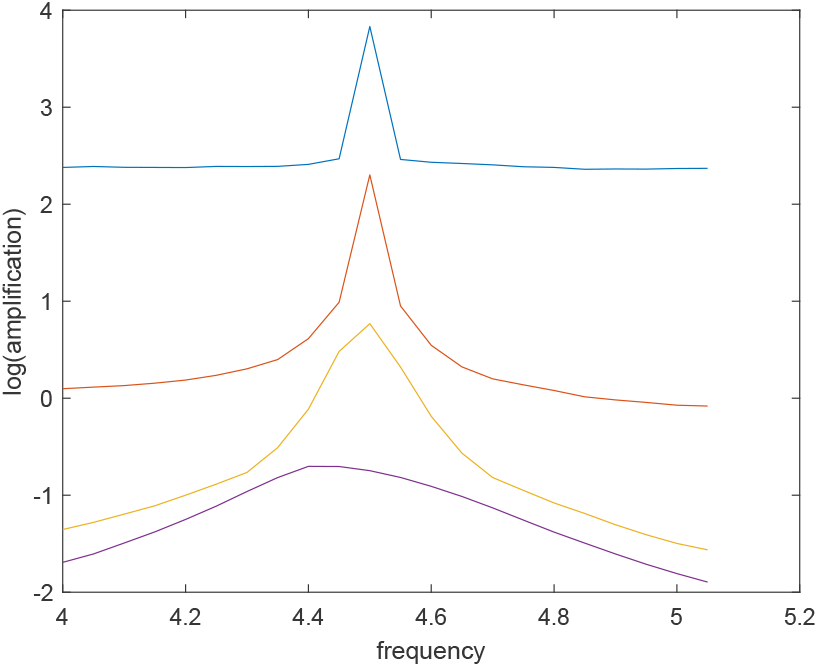
The amplitude amplification of the pure nonlinear oscillator for forced oscillations of the form *χ*(*t*) = *a* cos(*ωt*). The graphs are for four values of *a* : 10^−2^ (lowest graph), 10^−3^, 10^−4^ and 10^−5^ (highest). The parameters here are: *k*_*c*_ = 5, *k*_*d*_ = 40, *β*_*c*_ = 4.3, *β*_*d*_ = 0.3. The saturation parameter is *H* = 0.0065 and bifurcation parameter is Δ = 0.0282. The resonance frequency is *ω*_*c*_ = 4.5

## 4 Energy considerations for the full model (2.1) - (2.5)

In this section we derive apriori estimates for the solution of the system (2.1) - (2.5). For this purpose we multiply equations (2.2) by *h*_*t*_, and multiply equation (2.3) by *s*_*t*_. Adding these equations and arranging the different terms in a convenient form we obtain

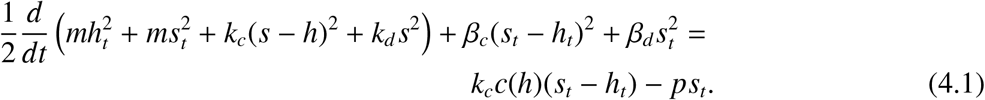

### Proposition 2

There exists a constant *C*_1_ such that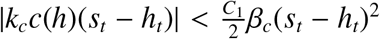. This inequality follows from a standard argument once we notice that |*c*(*h*)| is bounded.

We now integrate the identity (4.1) over 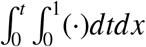, and argue:

### Proposition 3

Assume *p*(0, *t*) = *f* (*t*) is a bounded function. There exits a constant *C* such that the following inequality holds:

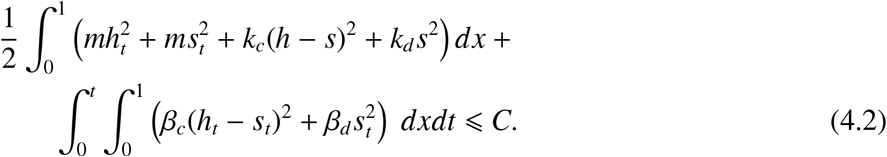

To derive inequality (4.2) we define a modified pressure 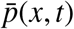 by writing

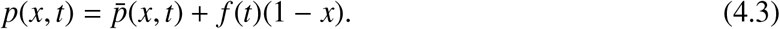

Clearly 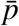 still satisfies equation (2.1), but now with homogeneous boundary conditions 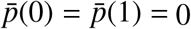. We then argue

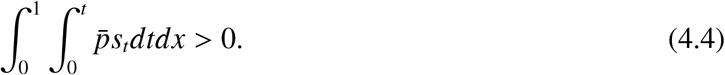

To prove this inequality we consider equation (2.1) for 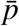 and *s*. Expand both 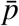 and *s* into Fourier series:

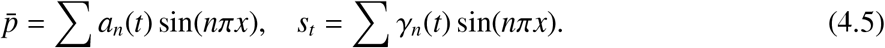

Substituting these expressions into equation (2.1) we obtain

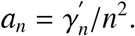

Therefore

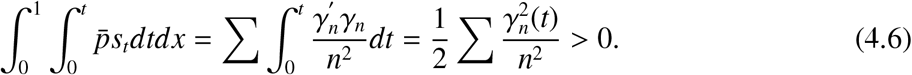

Using inequality (4.4), the fact that *f* (*t*)(1− *x*) is bounded, and Proposition 2 we obtain Proposition 3. Proposition 3 implies existence and regularity of the solution to the model equations (2.1)-(2.5).

Notice also the term (*s* − *h*)^2^ in equation (4.2) that hints at the closeness of *h* and *s*.

Denoting ∫_*per*_(·) *dt* integration over a period, then from equation (4.1) we conclude that ∫_*Per*_ *c*(*h*)*s*_*t*_*dt* and ∫_*Per*_ *ps*_*t*_*dt* are the work done on the BM by the OHC motility and by the applied pressure, respectively. Notice that the remaining term *c*(*h*)*h*_*t*_ on the right hand side on (4.1) does not contribute any work on the BM since it integrates over a period to zero.

## 5 Implementation of the model

In order to simulate the full model (2.1)-(2.5) we need to synchronize two independent families of frequencies, and furthermore to do so for each point *x* along the BM. One family is related to the critical frequencies of the pure oscillator that was studied in section 3. The other family is the natural frequencies of the BM, that vary along the BM according to the *place principle*. The first family is controlled by the nonlinear term *c*(*h*) and by the elastic and friction properties of the OHC and BM. It is determined by the imaginary part ℐ (*λ*(*x*)) of the bifurcating eigenvalue of the linearization matrix that was considered in section 3. The second family is controlled by the pressure difference across the BM and the BM and OHC elastic and friction coefficients. For our model to provide the needed amplification along the entire BM we need these two families to essentially agree (i.e. synchronize) at each point *x*. In addition to this synchronization, we need to control the real part ℛ (*λ*(*x*)) of the bifurcating eigenvalue. This function determines the growth of the instability at each point, and we want to keep it small (to be near the bifurcation point) and also essentially constant (so that all points along the BM are excited at the same rate). Another property that we would like to obtain, at least approximately, is that the place principle for the active cochlea is close to that principle in the passive cochlea.

Fulfilling these requirements is a difficult task that will be further discussed in the next section. Meanwhile we applied here an approximate approach, where we optimize the two degrees of freedom, i.e. the functions *k*_*c*_, *k*_*d*_, to achieve the requirements above up to a given tolerance. For this purpose we use equation (2.3) without the term *F*_*OHC*_, and the inequality *k*_*d*_ ≫ *β*_*d*_ to set 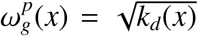 to be an approximation of the place principle for the passive BM. Similarly, we use the full equation (2.3), and inequality *k*_*c*_ ≫ *β*_*d*_ to define 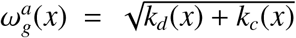. Next we define, in light of the Greenwood formula [9], for each point along a grid in the interval [0, 1] an empirical place principle function *ω*_*g*_(*x*) = 8.1 exp(−1.25*x*). The critical frequency distribution for the OHC is defined by *ω*_*c*_(*x*) := ℐ (*λ*(*x*)). The functions *k*_*c*_(*x*), *k*_*d*_(*x*) are then optimized, with fixed *β*_*c*_, *β*_*d*_, Δ, *H*, to meet the following conditions

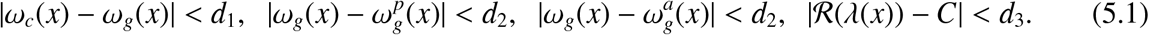

In our simulations we used the tolerance values *d*_1_ = 0.1, *d*_2_ = 0.3, *d*_3_ = 0.002 and the constant *C* = 0.002.

We thus obtained the functions *k*_*c*_(*x*), *k*_*d*_(*x*) as depicted in Figure 3. Notice that our model covers the frequencies interval (2.2, 8.1). The function *ω*_*c*_(*x*) is indeed quite close to the expected *ω*_*g*_(*x*), as demonstrated in Figure 4 where the circles are the points dictated by *ω*_*g*_ and the solid graph is the critical frequency distribution *ω*_*c*_(*x*) for each of the pure oscillators. We emphasize that our selection of parameters is somewhat arbitrary and is not meant to represent a specific real-world cochlea. It is done only in order to demonstrate the model’s implications.

**Figure 3:**
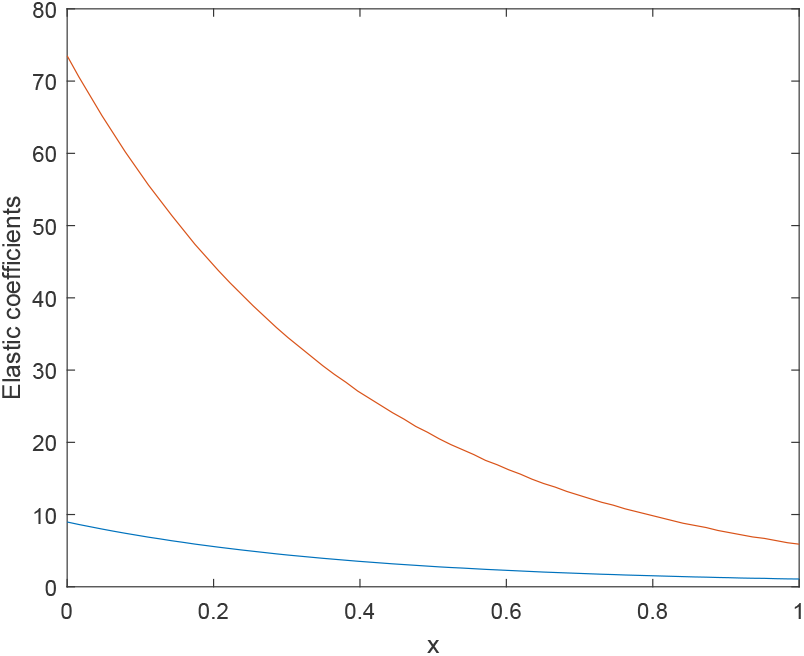
The elastic functions *k*_*c*_(*x*) (lower curve) and *k*_*d*_(*x*) (upper curve).

**Figure 4:**
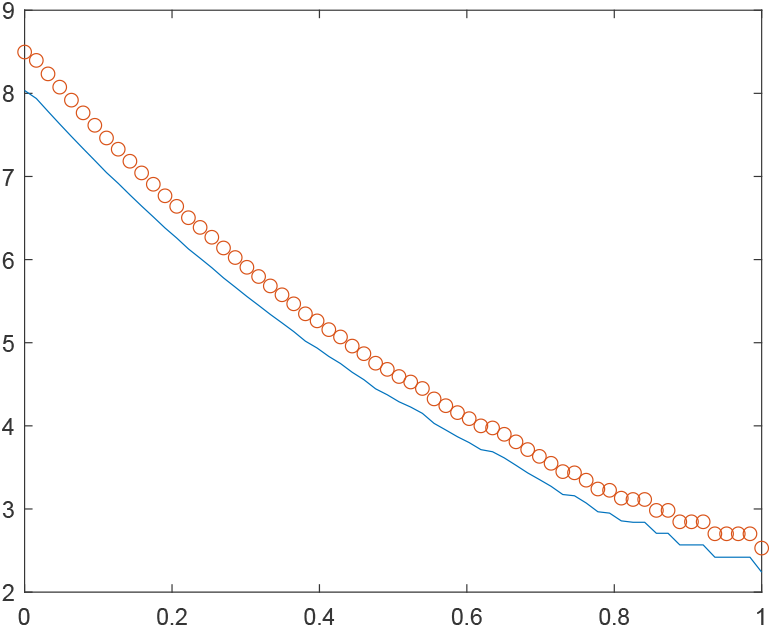
The critical frequency distribution *ω*_*c*_(*x*) (solid curve) vs. the place principle frequency distribution *ω*_*g*_(*x*) (circles).

Using the functions *k*_*c*_(*x*), *k*_*d*_(*x*) selected above, and using the parameters *β*_*c*_ = 1.1, *β*_*d*_ = 0.75, Δ = 0.1, *H* = 0.0065, we can run simulations of the full model. We used second order numerical schemes in time and space. Space was discretized by 64 points and the time step was *dt* = 0.01. We set the boundary condition *p*(0, *t*) := *P* = 10^−5^(cos(5.2*t*) + cos(3.8*t*) + cos(3.6*t*)). In Figure 5 we draw for each point *x* along the BM the function *S* (*x*) = max_*t*_ *s*(*x, t*). The response curve easily identifies three sharp amplified peaks, precisely at the expected location according to *ω*_*c*_(*x*).

**Figure 5:**
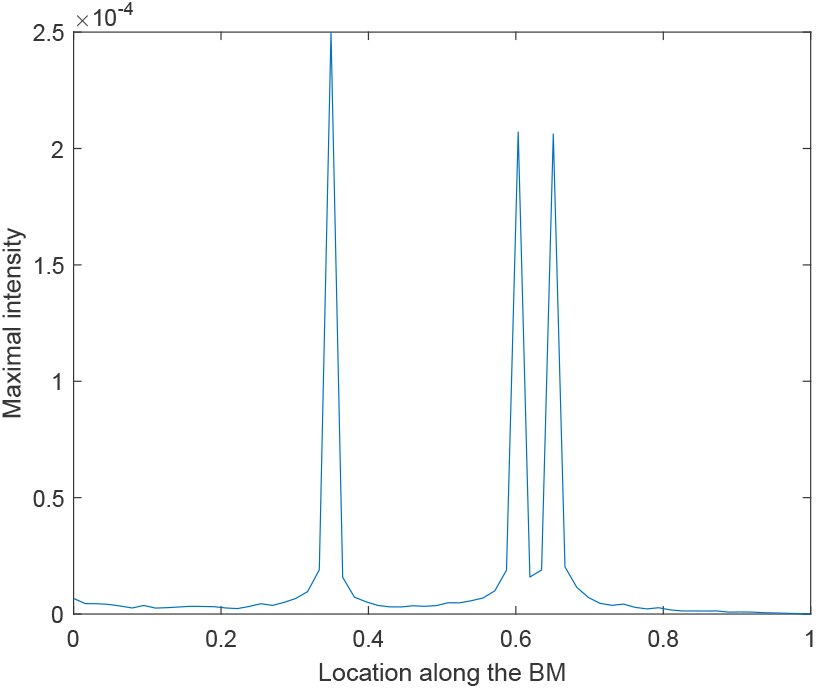
Simulation of the full model (equations (2.1)-(2.5)). The system is driven by the pressure difference *p*(0, *t*) = 10^−5^(cos(5.24*t*)+cos(3.8*t*)+cos(3.6*t*)). The friction parameters are *β*_*c*_ = 1.1, *β*_*d*_ = 0.75. The saturation parameter is *H* = 0.0065 and the control parameter is Δ = 0.1. The elastic coefficients *k*_*c*_(*x*), *k*_*d*_(*x*) are depicted in Figure 3. The figure depicts the maximal (over many periods) intensity of the deflection *s*(*x, t*) of the BM at each point *x* along the BM.

It is interesting to compare the BM amplitude *S* (*x*) above with a similar curve for a *passive* BM model that consists of equations (2.1) and (2.3) without the term *F*_*OHC*_. The simulation of this model with the same initial condition *p*(0, *t*) as in the previous example is drawn in Figure 6. Here we used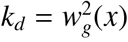, and we set *β*_*d*_ = 0.1. As seen in Figure 6, the passive BM model gives rise to broad peaks with very small amplification. Notice that although the friction *β*_*d*_ here is much smaller than in the full model (e.g. Figure 5), the passive BM can barely distinguish between the frequencies 3.8 and 3.6, in sharp contrast to the clear distinction shown in Figure 5.

**Figure 6:**
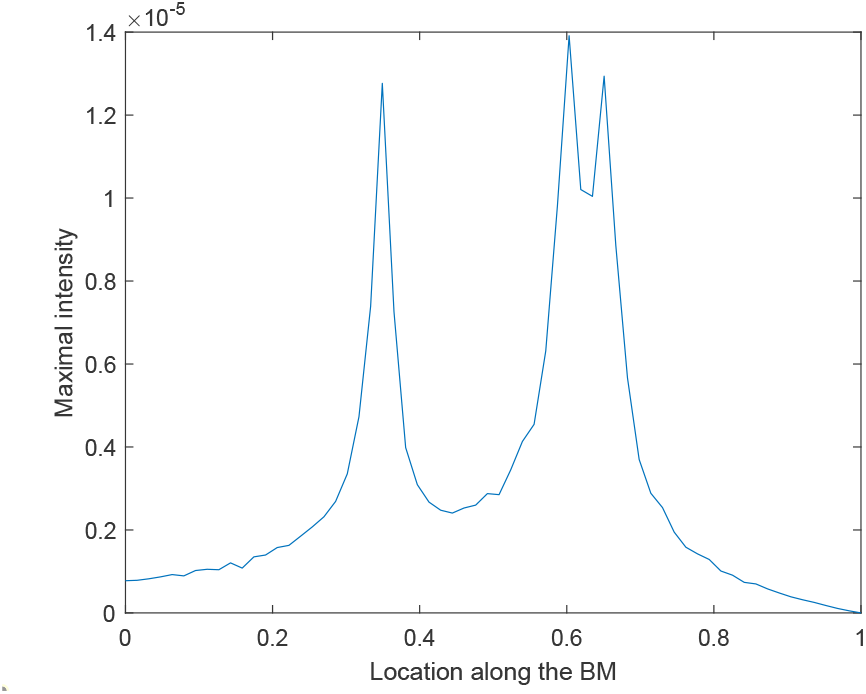
Simulation of the passive cochlea (equations (2.1), (2.3) without the term *F*_*OHC*_). The system is driven by the pressure difference *p*(0, *t*) = 10^−5^(cos(5.24*t*) + cos(3.8*t*) + cos(3.6*t*)). The friction term is *β*_*d*_ = 0.1. The elastic coefficient here is 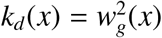. The figure depicts the maximal (over time) intensity of the deflection *s*(*x, t*) of the BM at each point *x* along the BM for many periods.

We now simulate the performance of the full model under noisy data. In the next simulation the input consisted of noisy data in the interval 0 < *t* < 400 and the same input as in Figure 5 imposed over the noisy data in the time interval 400 < *t* < 800. The noisy signal is modeled as 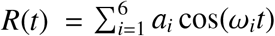, where the frequencies *ω*_*i*_ are uniformly distributed over (2, 8) and the amplitudes are uniformly distributed over 3 · 10^−5^[0, 1]. The random variables are switched every 100 time steps, where the entire interval is divided into 80, 000 time steps. The input is depicted in Figure 7. Since the noise amplitude is larger than the pure signal *P*, it is hard to discern the change between the first and second time intervals. The maximal amplitude *S* (*x*) of the BM over many periods is depicted in Figure 8. The + symbols stand for *S* (*x*) in the interval (0, 400), while the solid line is for *S* (*x*) in the time interval (400, 800).

**Figure 7:**
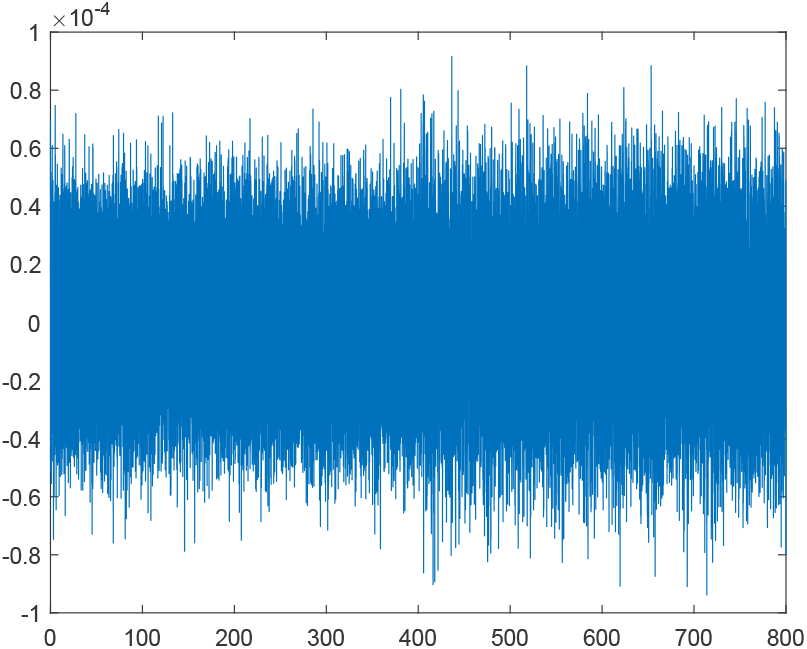
Noisy signal during 0 < *t* < 400 and input consisting of noisy signal plus the sum of three pure frequencies in the interval 400 < *t* < 800.

**Figure 8:**
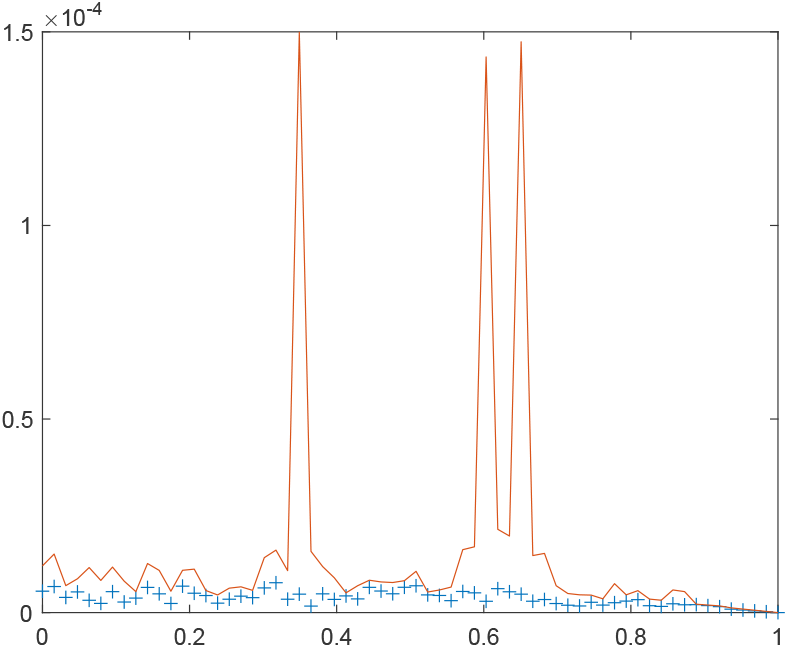
Simulation of the full model (equations (2.1)-(2.5)) for the input depicted in Figure 7. The input consisted of noisy data only in the interval 0 < *t* < 400. In the interval 400 < *t* < 800 the input consisted of similar noisy data plus three pure tones 10^−5^(cos(5.24*t*) + cos(3.8*t*) + cos(3.6*t*)). The figure depicts the maximal (over time) intensity *S* (*x*) of the deflection *s*(*x, t*) of the BM at each point *x* along the BM for many periods in the interval 0 < *t* < 400 (marked by + symbol) and also *S* (*x*) in the interval 400 < *t* < 800 (solid line).

## 6 Discussion

We presented a new model for the mutual interaction of the OHC and the BM, leading to many desired properties, such as amplification and saturation of the incoming audio signal, sharpening the spectral decomposition of the signal and dramatic filtering of noise. The key component of the model is the motility of the OHC, induced by inclination of the OHB, leading to a dynamical system that undergoes a Hopf bifurcation under proper selection of different parameters.

The effect of the specific type of OHC motility assumed here, on the signal processing of a slice of the OoC, was already established in an earlier paper [3]. A novel feature in the current paper is the synchronization of two independent families of oscillators that enabled us to combine the OHC motility model with the BM vibration initiated by the pressure difference across it. The first family is given by a set of critical frequencies determined by the nonlinear motility of the OHC and its interaction with the BM elasticity. The second family is the classical oscillator determined by the elastic response of the BM to the pressure difference between the scala media and scala tympani. This oscillator gives rise to the *place principle*, where each point along the BM is associated with a specific frequency. It forms the first step towards the decomposition of sound to its spectral components. In order to synchronize these two families we optimized the elastic functions *k*_*c*_(*x*) of the OHC and *k*_*d*_(*x*) of the BM so that the two frequency families would be essentially the same at each point *x* along the entire BM. At the same time we imposed on *k*_*c*_, *k*_*d*_ an additional condition, namely, that the real part of the bifurcation eigenvalue would be approximately constant along the BM, in order to ensure that the instability develops at similar rates at each point.

This synchronization enabled us to simulate the BM-OHC system as a whole and to show that the combined model indeed gives rise to the cochlea properties listed above. In particular we showed that the new model implies sharp spectral decomposition and amplification (Figure 5), and noise filtering (Figure 8). Since the pure oscillator model is at the core of the model here and in [3], we studied it in more detail in section 3. In particular we demonstrated the unusual role played by the OHC friction. An important feature of our model is that it shows explicitly that the OHC does actual work on the BM, in addition to the work performed by the force exerted on it by the pressure difference across the BM [21].

The model presented here was developed under a number of simplifying assumptions. It can be expanded in different ways. One challenge is to combine the horizontal model here with the vertical model in [3]; namely, to connect the 1d BM approximation with a 1d OoC model that includes the DC, the OHB, and the IHB. Another extension is towards 2d or 3d models for the fluid flow in the scala media and scala tympani, and similarly for the fluid flow in the subtectorial endolymph. Recent work [14] and [10] argued for the importance of going beyond 1d model for the fluid equations. Another important aspect to consider is a more realistic space-dependent model for the friction and mass density parameters that we took here as constant.

The search for more realistic elastic and friction coefficients is related to the requirements, mentioned in section 5, on the problem parameters, namely, synchronization of different types of frequencies and controlling the instability rate. An alternative approach to the method of section 5 is to start from an *experimental* distribution *ω*_*g*_(*x*) for the place principle, obtained say, for a passive cochlea. Assume that *ω*_*g*_ is very close to that of the active cochlea, and then select *k*_*c*_, *k*_*d*_ (and maybe other parameters) to meet the first and fourth conditions of equation (4.6). Another method, more theoretical in nature, is to write down a WKB (e.g. [16], [17]) expansion for linear regime the full model (2.1)-(2.5). Such an expansion was used for the passive BM, with varying elastic and friction coefficients, to find the location *x*_*g*_(*ω*) along the BM where the its deflection is maximal for a given frequency. Writing such an expansion for the full model should give 3 *simultaneous* equations for ℐ (λ(*x*)), ℛ (λ(*x*)) and 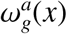 involving all the model parameters.

Finally, our nonlinear motility function *c*(*h*) is symmetric about the origin and is saturated using the tanh function. This is a common model in neural networks. However, it would interesting to test alternative saturation models.

## 7 Appendix: The fluid-BM 1d model

Consider a straightened-out two-chamber cochlea model where the two chambers are separated by the BM. The length of the chamber is *L*, and the height and width are *l*. The fluid equation in the chamber ± is

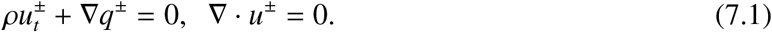

Here *u*^±^ and *q*^±^ are the velocity and pressure, respectively, in each chamber, and ρ is the fluid density. Equations (7.1) can be written as a single equation 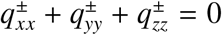 for the pressure. We integrate this equation over a *y* − *z* cross section of each chamber. Assuming that since *l* ≪ *L*, then to leading order 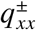 is independent of *y, z*, we obtain

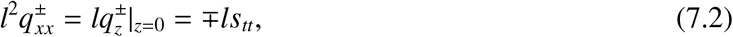

where we used equation (7.1) and continuity of the acceleration of the BM-fluid interface.

The BM force equation is *ms*_*tt*_ + *k*_*d*_ *s* + *b*_*d*_ *s*_*t*_ = *F*_*OHC*_− *l*[*q*], where [*q*] = *q*^+^(*z* = 0) − *q*^−^(*z* = 0) is the pressure difference across the BM. Defining *l*[*q*] := *p*, and scaling the *x* coordinate by *L* (while retaining the *x* notation for the scaled coordinate) we obtain the following set of equations

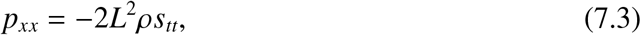

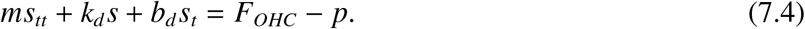

Finally using *L* = 3 · 10^−2^ *m*^2^ and ρ = 10^3^ *Kg*/*m*^3^, we obtain 2*L*^2^ρ = *O*(1). Since there are already a number of free parameters in the model, we replace this term with 1.

